# Subject, session and task effects on power, connectivity and network centrality: a source-based EEG study

**DOI:** 10.1101/673343

**Authors:** Sara M. Pani, Marta Ciuffi, Matteo Demuru, Giovanni Bazzano, Ernesto D’aloja, Matteo Fraschini

**Affiliations:** Department of Biomedical Sciences, PhD Program in Neuroscience, University of Cagliari, I-09042, Italy; Department of Medical Sciences and Public Health - Forensic Science Unit - University of Cagliari, Cagliari, Italy; Department of Electrical and Electronic Engineering, University of Cagliari, Cagliari, I-09123, Italy

**Keywords:** EEG, individuality, connectivity, task-switching

## Abstract

Inter-subjects’ variability in functional brain networks has been extensively investigated in the last few years. In this context, unveiling subject-specific characteristics of EEG features may play an important role for both clinical (e.g., biomarkers) and bio-engineering purposes (e.g., biometric systems and brain computer interfaces). Nevertheless, the effects induced by multi-sessions and task-switching are not completely understood and considered. In this work, we aimed to investigate how the variability due to subject, session and task affects EEG power, connectivity and network features estimated using source-reconstructed EEG time-series. Our results point out a remarkable ability to identify subject-specific EEG traits within a given task together with striking independence from the session. The results also show a relevant effect of task-switching, which is comparable to individual variability. This study suggests that power and connectivity EEG features may be adequate to detect stable (over-time) individual properties within predefined and controlled tasks.

## I. INTRODUCTION

Despite most neuroimaging studies still tend to treat human brain features as stable and homogeneous characteristics within a group, it is important to highlight that, in contrast, individual variability may play a relevant role in this context [1], [2]. The way in which each brain is unique and could be distinguished amidst a myriad of other brains is fascinating, but unveiling the underlying subject-specific characteristics is crucial for both clinical (e.g., biomarkers) and bio-engineering purposes (e.g., biometric systems and brain computer interfaces). Recent studies have already highlighted the implications of individual variation for personalized approaches to mental illness [3], ADHD [4] and in the developing brain [5]. It has been also reported that these functional traits are familial, heritable and stable over a long time interval [6], [7]. Electroencephalographic (EEG) time-frequency [8] and connectivity-based [9], [10] features have shown subject-specific characteristics comparable in terms of performance to other more common fingerprints. Nevertheless, the performance of EEG-based biometric systems seems to be not independent from the specific connectivity metric, scarcely investigated in terms of permanence and tend to decrease in a between-tasks scenario [11]. From this new perspective, with the clear evidence that functional brain networks vary across individuals, few studies investigated to what extent these subject-specific traits are stable over time and over different states. Using functional Magnetic Resonance Imaging (fMRI), Gratton et al. [12] reported that functional networks are suited to detect stable individual characteristics with a limited contribution from task-state and day-to-day variability, thus suggesting their possible utility in the personalized medicine approach. Similarly, Cox et al. [13], using EEG scalp level analysis, have reported that, despite a shared structure is still discernible across individuals, well-defined subject-specific and stable over-time network profiles were clearly detectable. In this study we aim to investigate if these subject-specific traits are still detectable, stable over time and consistent among different tasks using an EEG source level approach. This approach should provide a more accurate description of the underlying network [14] since the connectivity estimates should be less prone to volume conduction and signal leakage problems. In order to investigate this question we analyzed source-reconstructed EEG time-series using three different and widely used analyses: Power Spectral Density (PSD), Phase Locking Value (PLV) [15] and nodal centrality network approaches, namely Eigenvector Centrality (EC). PSD has been shown to capture relevant subject-specific information [8] and represents one of the more simple and interpretable EEG features. PLV, in combination with weighted Minimum Norm Estimator (wMNE) [16], provides a good estimate of the functional brain organization in EEG [17] and, despite the PLV is not completely independent from the PSD [18], is known to be affected by volume conduction and signal leakage, it still performs better than other common connectivity metrics in terms of subject authentication [11]. Moreover, as previously stated, the PLV was recently used at scalp-level to investigate variability and stability of large-scale cortical oscillation patterns [13]. Finally, it was reported that the EC, which captures more information about the network topology then straightforward measure such as the degree, represents a promising measure to design of EEG-based biometric systems [9]. The analysis was performed on a novel EEG dataset consisting of fourteen healthy subjects, recorded over two different sessions (after four weeks) and performing four different tasks. All the code is freely available in a Github repository at the following link: https://github.com/matteogithub/individuality.

## II. METHOD

### A. EEG PREPROCESSING

All the preprocessing steps were performed using the freely available toolbox EEGLAB (version 13_6_5b) [19]. The raw EEG signals were re-reference to common average reference and band-pass filtered (with fir1 filter type) between 1 and 70 Hz and a notch filter set to 50 Hz was also applied. All the recordings were visually inspected and segments with clear artifacts were rejected and not further analyzed.

### B. SOURCE RECONSTRUCTION

In order to obtain the source-reconstructed time-series, the Brainstorm software (version 3.4) [20] was used to compute the head model with a symmetric boundary element method in Open-MEEG [21] based on the anatomy derived from the ICBM152 brain. EEG time-series at source level were reconstructed using whitened and depth-weighted linear L2 minimum norm estimate (wMNE) [16], [22] and projected onto 68 regions of interest (ROIs) as defined by the Desikan-Killiany atlas [23].

### C. FEATURES EXTRACTION

After the EEG time-series were reconstructed at source level, in order to increase the quality of the analysis, for each subject, each task and each session, we selected the best (less contaminated) 10 EEG epochs (segments of 5 seconds) ordering all the available epochs on the basis of the three-sigma rule (consequently discarding segments presenting values over than 3 standard deviations from the mean) [24]. Successively, for each selected epoch we have extracted three different features vectors, respectively for PSD, PLV and EC, representing the individual profiles or subject fingerprints. For the PSD analysis, the features vector, for each single epoch, was composed of the 272 entries representing the relative power (extracted using the Welch method) of four frequency bands (delta [1 - 4 Hz], theta [4 - 8 Hz], alpha [8 - 13 Hz] and beta [13 - 30 Hz]), separately for each of the 68 regions of interest. For the PLV analysis, the features vector, for each single epoch and for each frequency band, was composed of 2.278 entries representing the connectivity profile (upper triangular of the connectivity matrix), where each entry was computed as:

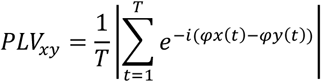

where T is the epoch length and φ is the instantaneous phase. For the network analysis, in order to keep a nodal resolution, we have computed the EC, a centrality measure based on the spectral decomposition of the weighted connectivity matrix [25]. In this latter case the features vector, for each single epoch and separately for each frequency band, was composed of 68 entries, each representing the centrality value of the corresponding ROI. As a final step, in order to estimate the similarity among each pairs of possible observations (between-epochs), we computed the Euclidian distance between features vectors (individual profiles) independently for PSD, PLV and EC analysis, thus obtaining, for each analysis, a square and symmetric matrix of distances, with the dimension equals to (number of subjects) * (number of sessions) * (number of tasks) * (number of epochs) as shown in Figure 1. From this distances matrix, we have computed the average distances across epochs for each of the following six scenarios: (i) within-task, within-session and within-subject; (ii) between-tasks, within-session and within-subject; (iii) between-sessions, within-task and within-subject; (iv) between-sessions, between-tasks and within-subject; (v) within-task, within-session and between-subjects; (vi) all-between. All the code, developed in Matlab, reporting the extraction of the profiles and their comparison, is freely available at the following link in Github: https://github.com/matteogithub/individuality.

**FIGURE 1.**
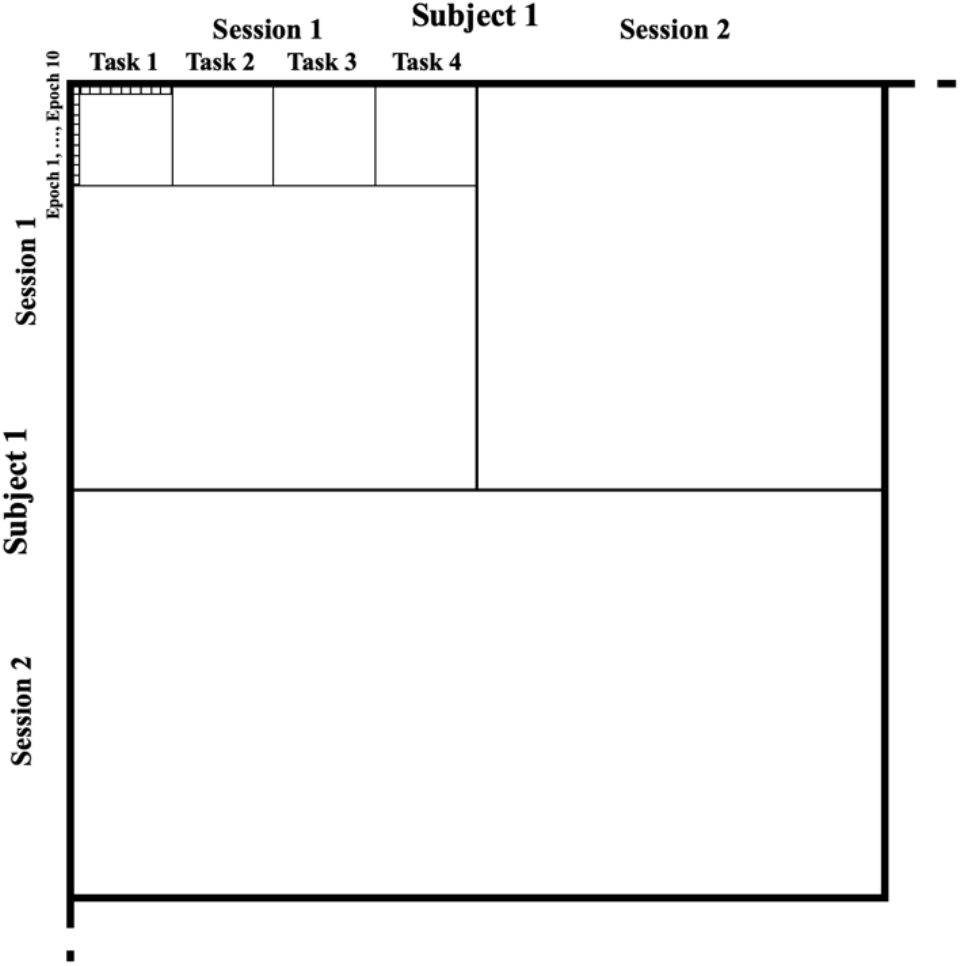
A schematic representation of the first block (one subject) of the matrix containing the distances. The main diagonal contains zeros.

### D. STATISTICAL ANALYSIS

The statistical analysis was performed by using the non-parametric Kruskal-Wallis test followed by two-stage linear step-up procedure of Benjamini, Krieger and Yekutieli [26] to account for the multiple comparison problem.

## III. DATASET

Fifteen healthy volunteers (7 females, mean age 31.9 ± 3.1 years, range 28 – 38) were enrolled in the present study. Informed consent was obtained prior to the recordings and the study was approved by the local ethics committee. EEG signals were recorded using a 61 channels EEG system (Brain QuickSystem, Micromed, Italy) during four different tasks and repeated over two different session (the second acquired four weeks later from the first). Recordings were acquired in a sitting position in a normal daylight room; a dimly lit and sound attenuated room and supine position were avoided to prevent drowsiness. Signals were digitized with a sampling frequency of 1024 Hz with the reference electrode placed in close approximation of the electrode POz. The four tasks consisted of: (i) five minutes eyes-closed resting-state, (ii) five minutes eyes-open resting-state, (iii) two minutes eyes-closed simple mathematical task and (iv) two minutes eyes-closed complex mathematical task. During the simple mathematical task, the subjects were asked to perform multiple subtractions, while during the complex mathematical task, subjects were asked to perform a series two digits multiplications. Three subjects were excluded from the analysis due to low quality of the EEG recordings and another one missed the second session.

## IV. RESULTS AND DISUSSIONS

### A. PSD

Results derived from PSD analysis are shown in Figure 2 and the corresponding statistics are summarized in Table 1. The lower distances were observed for the within-task, within-session, within-subject scenario (0.95 ± 0.34) and for the between-sessions, within-task, within-subject scenario (0.87 ± 0.30). The distances increased for the between-tasks scenarios, both for within-session (3.06 ± 1.20) and for between-sessions (3.09 ± 1.19). The distances further increased for the between-subjects’ scenarios, both for within-session, within-task (3.23 ± 1.00) and for all between (3.59 ± 1.18).

**FIGURE 2.**
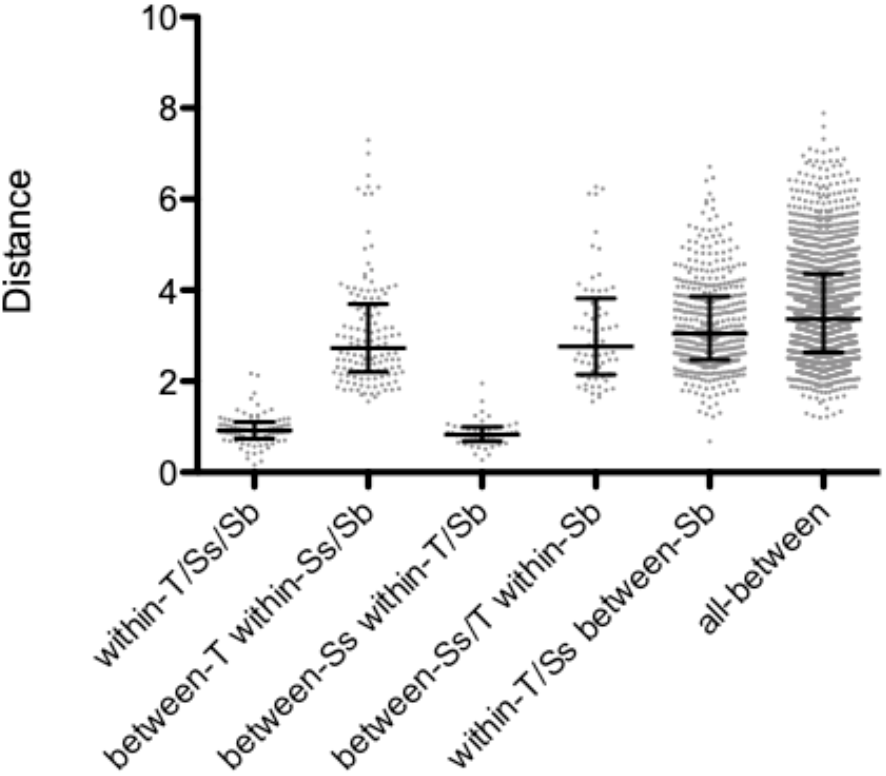
Scatterplot of distances obtained by using the PSD approach. Bars represent median and interquartile range. T is for task, Ss for session and Sb for subject.

**TABLE I.** Statistical results for PSD Analysis

| STATISTICAL RESULTS FOR PSD ANALYSIS |  |  |
| --- | --- | --- |
|  | Mean rank diff. | p-value |
| w-T/Ss/Sb vs. b-T w-Ss/Sb | -873.129 | <0.0001 |
| w-T/Ss/Sb vs. b-Ss w-T/Sb | 14.6023 | 0.9056 |
| w-T/Ss/Sb vs. b-Ss/T w-Sb | -907.163 | <0.0001 |
| w-T/Ss/Sb vs. w-T/Ss b-Sb | -1033.61 | <0.0001 |
| w-T/Ss/Sb vs. all-b | -1209.25 | <0.0001 |
| b-T w-Ss/Sb vs. b-Ss w-T/Sb | 887.731 | <0.0001 |
| b-T w-Ss/Sb vs. b-Ss/T w-Sb | -34.0341 | 0.7350 |
| b-T w-Ss/Sb vs. w-T/Ss b-Sb | -160.484 | 0.0153 |
| b-T w-Ss/Sb vs. all-b | -336.122 | <0.0001 |
| b-Ss w-T/Sb vs. b-Ss/T w-Sb | -921.765 | <0.0001 |
| b-Ss w-T/Sb vs. w-T/Ss b-Sb | -1048.21 | <0.0001 |
| b-Ss w-T/Sb vs. all-b | -1223.85 | <0.0001 |
| b-Ss/T w-Sb vs. w-T/Ss b-Sb | -126.450 | 0.1509 |
| b-Ss/T w-Sb vs. all-b | -302.088 | 0.0003 |
| w-T/Ss b-Sb vs. all-b | -175.638 | <0.0001 |

Statistics refer to non-parametric multiple comparison tests (on mean rank) using the FDR correction approach, where w- and b- refers to within and between scenarios respectively. T, Ss and Sb refer to task, session and subject.

### B. CONNECTIVITY

Results derived from PLV based analysis in the beta band are consistent with those obtained by PSD as shown in Figure 3 and the corresponding statistics summarized in Table 2. Again, the lower distances were observed for the within-task, within-session, within within-subject scenario (4.82 ± 0.33) and for the between-sessions, within-task, within-subject scenario (4.40 ± 0.34). The distances increased for the between-tasks scenarios, both for within-session (5.74 ± 0.85) and for between-sessions (5.85 ± 0.96). Finally, the distances further increased for the between-subjects’ scenarios, both for within-session, within-task (7.07 ± 0.76) and for all between (7.25 ± 0.87).

**FIGURE 3.**
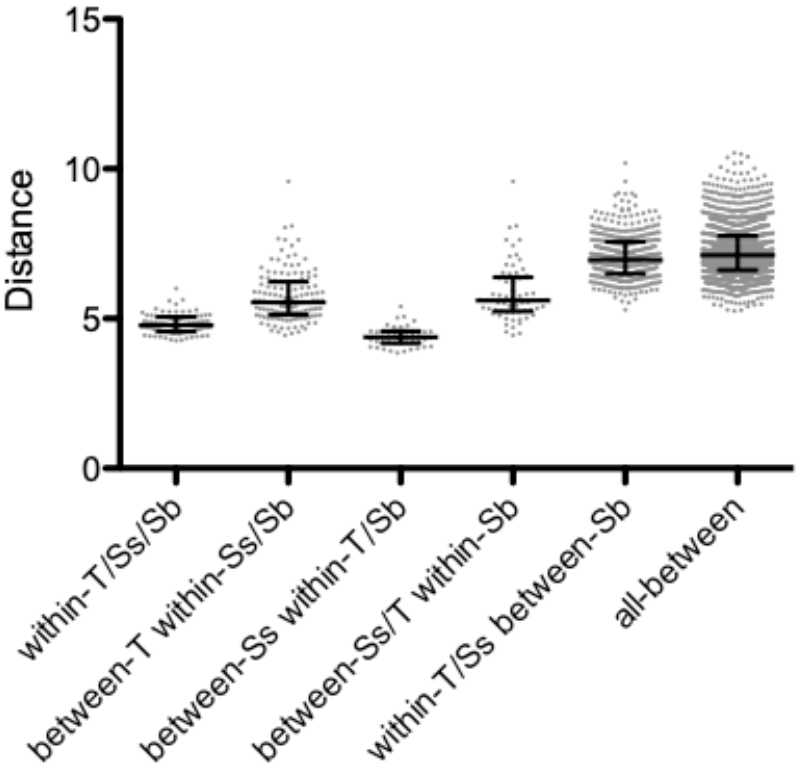
Scatterplot of beta band distances obtained by using the PLV connectivity approach. Bars represent median and interquartile range. T is for task, Ss for session and Sb for subject.

**TABLE II.** Statistical results for PLV Beta band

|  | Mean rank diff. | p-value |
| --- | --- | --- |
| w-T/Ss/Sb vs. b-T w-Ss/Sb | -317.8 | 0.0005 |
| w-T/Ss/Sb vs. b-Ss w-T/Sb | 61.03 | 0.6202 |
| w-T/Ss/Sb vs. b-Ss/T w-Sb | -382.5 | 0.0004 |
| w-T/Ss/Sb vs. w-T/Ss b-Sb | -1104 | <0.0001 |
| w-T/Ss/Sb vs. all-b | -1217 | <0.0001 |
| b-T w-Ss/Sb vs. b-Ss w-T/Sb | 378.8 | 0.0011 |
| b-T w-Ss/Sb vs. b-Ss/T w-Sb | -64.72 | 0.5198 |
| b-T w-Ss/Sb vs. w-T/Ss b-Sb | -786.1 | <0.0001 |
| b-T w-Ss/Sb vs. all-b | -899.3 | <0.0001 |
| b-Ss w-T/Sb vs. b-Ss/T w-Sb | -443.6 | 0.0006 |
| b-Ss w-T/Sb vs. w-T/Ss b-Sb | -1165 | <0.0001 |
| b-Ss w-T/Sb vs. all-b | -1278 | <0.0001 |
| b-Ss/T w-Sb vs. w-T/Ss b-Sb | -721.4 | <0.0001 |
| b-Ss/T w-Sb vs. all-b | -834.6 | <0.0001 |
| w-T/Ss b-Sb vs. all-b | -113.1 | 0.0017 |

Statistics refer to non-parametric multiple comparison tests (on mean rank) using the FDR correction approach, where w- and b- refers to within and between scenarios respectively. T, Ss and Sb refer to task, session and subject.

The results show a similar pattern, still slightly less marked, also for the alpha band as shown in Figure 4 and the corresponding statistics summarized in Table 3. In this case, again the lower distances were observed for the within-task, within-session, within-subject scenario (8.25 ± 0.86) and for the between-sessions, within-task, within-subject scenario (7.55 ± 0.83). The distances increased for between-tasks scenarios, both for within-session (9.21 ± 1.26) and for between-sessions (9.32 ± 1.27). Finally, the distances further increased for the between-subjects’ scenarios, both for within-session, within-task (10.86 ± 2.11) and for all between (11.01 ± 2.16).

**FIGURE 4.**
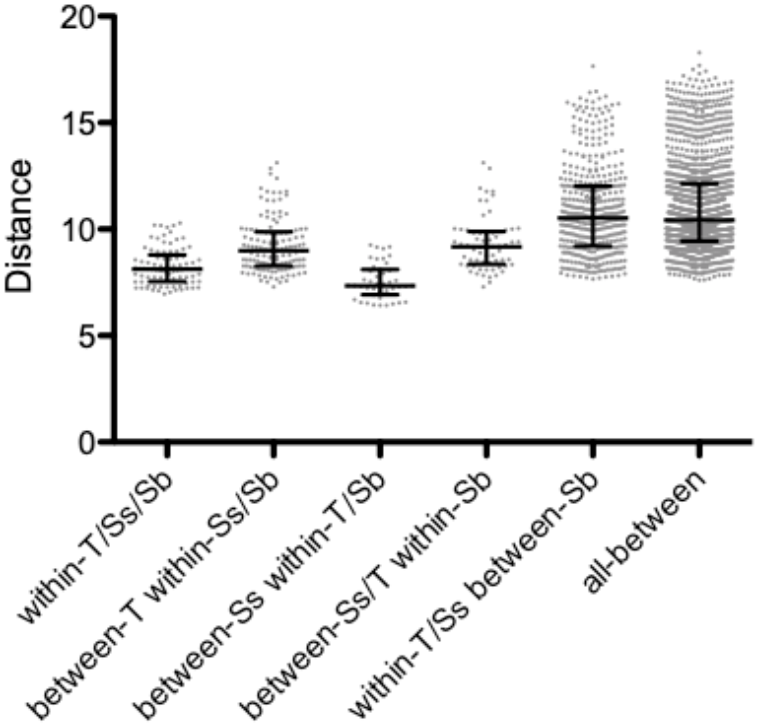
Scatterplot of alpha band distances obtained by using the PLV connectivity approach. Bars represent median and interquartile range. T is for task, Ss for session and Sb for subject.

**TABLE III.**
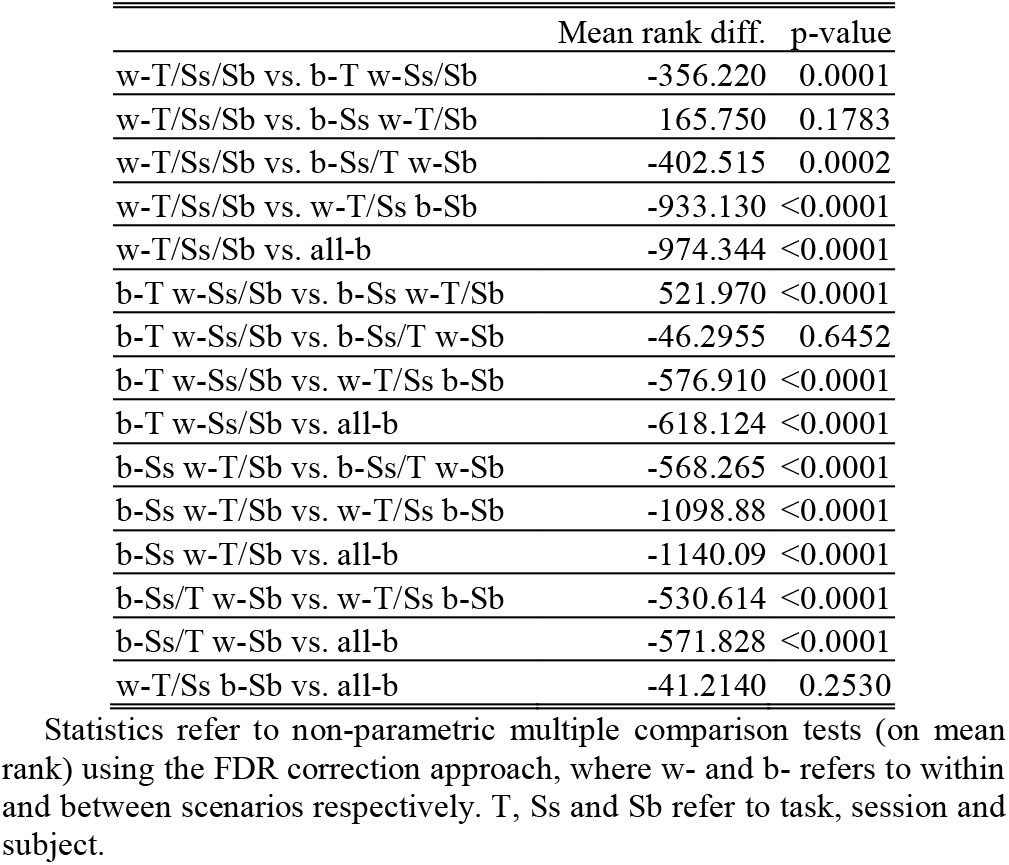
Statistical results for PLV Alpha band

### C. NETWORK CENTRALITY

Results derived from the application of EC on PLV based analysis (in the beta band) are still consistent with the previously reports, as shown in Figure 5 and the corresponding statistics summarized in Table 4. Again, the lower distances were observed for the within-task, within-session, within-subject scenario (0.12 ± 0.01) and for the between-sessions, within-task, within-subject scenario (0.11 ± 0.01). The distances increased for the between-tasks scenarios, both for within-session (0.14 ± 0.02) and for between-sessions (0.14 ± 0.02). Finally, the distances further increased for the between-subjects’ scenarios, both for within-session, within-task (0.17 ± 0.03) and for all between (0.17 ± 0.03).

**FIGURE 5.**
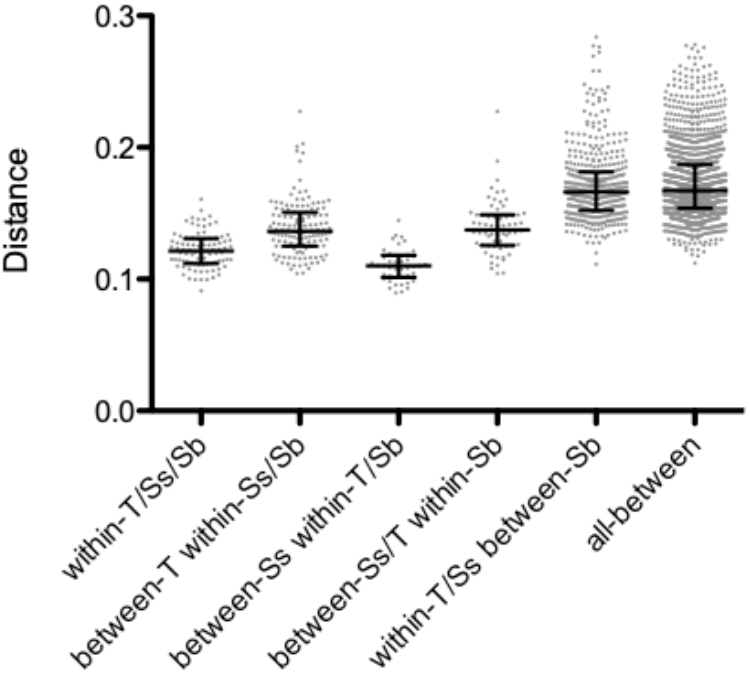
Scatterplot of alpha band distances obtained by using the PLV connectivity approach and eigenvector centrality. Bars represent median and interquartile range. T is for task, Ss for session and Sb for subject.

**TABLE IV.** Statistical results for Eigenvector Centrality

| TABLE IV<br>STATISTICAL RESULTS FOR EIGENVECTOR CENTRALITY |  |  |
| --- | --- | --- |
|  | Mean rank diff. | p-value |
| w-T/Ss/Sb vs. b-T w-Ss/Sb | -309.489 | 0.0007 |
| w-T/Ss/Sb vs. b-Ss w-T/Sb | 93.5795 | 0.4473 |
| w-T/Ss/Sb vs. b-Ss/T w-Sb | -307.186 | 0.0047 |
| w-T/Ss/Sb vs. w-T/Ss b-Sb | -1084.05 | <0.0001 |
| w-T/Ss/Sb vs. all-b | -1134.76 | <0.0001 |
| b-T w-Ss/Sb vs. b-Ss w-T/Sb | 403.068 | 0.0005 |
| b-T w-Ss/Sb vs. b-Ss/T w-Sb | 2.30303 | 0.9817 |
| b-T w-Ss/Sb vs. w-T/Ss b-Sb | -774.561 | <0.0001 |
| b-T w-Ss/Sb vs. all-b | -825.269 | <0.0001 |
| b-Ss w-T/Sb vs. b-Ss/T w-Sb | -400.765 | 0.0020 |
| b-Ss w-T/Sb vs. w-T/Ss b-Sb | -1177.63 | <0.0001 |
| b-Ss w-T/Sb vs. all-b | -1228.34 | <0.0001 |
| b-Ss/T w-Sb vs. w-T/Ss b-Sb | -776.864 | <0.0001 |
| b-Ss/T w-Sb vs. all-b | -827.573 | <0.0001 |
| w-T/Ss b-Sb vs. all-b | -50.7081 | 0.1596 |

Statistics refer to non-parametric multiple comparison tests (on mean rank) using the FDR correction approach, where w- and b- refers to within and between scenarios respectively. T, Ss and Sb refer to task, session and subject.

### D. DISCUSSIONS

In summary, in this work we aimed to investigate how the variability due to subject, session and task affects EEG power, connectivity and network features estimated using source-reconstructed EEG time-series. Despite this question was extensively investigated using fMRI [2], [6], [12], high density EEG, which still represents a very important and useful clinical tool, have received less attention in this context. Although, numerous studies have investigated the possibility to use EEG signals to develop biometric systems, only recently more attention was devoted to the study of subject variability and stability over-time and states [13]. The results of this study show three main relevant points. First, as expected, for all the different analyses, PSD, PLV and EC based approaches, the lower distances were observed in the scenario corresponding to a simple between epochs scheme, within the same subject, the same session and the same task. It should be highlighted that this also represents the more common scenario in which studies do not consider the variance induced by subject-specific traits, multi-sessions and/or by multi-tasks setup. Second, probably the more interesting finding, the distances obtained using the between-sessions, within-task, within-subject scenario are comparable with the previous one (namely, within-session scenario) for all the performed analyses. This finding clearly indicates that the variance due to the session is therefore negligible. Third, conversely, the effect due to the task (task-switching) is substantial, as also highlighted by the statistics and consistent for all the different analyses (i.e., PSD, PLV in beta and alpha bands and EC analysis). Moreover, the second point is further confirmed by the between-sessions and between-tasks scenario, where again it is still evident the low effect due to the session switching. Finally, as expected, the distances strongly increase in the between-subjects scenario, showing a clear effect due to specific subject, thus confirming the importance to address the issue related with the variance within a group. The reported results support, as recently reported using a scalp-level EEG analysis [13], the existence of well-defined subject-specific profiles and that these features may be considered stable over a defined and limited time range. These results are also in line with the fact that task-invariant subject-specific features are stronger than task-dependent group profiles. Finally, the reported findings also represent an important confirmation obtained at source level, of the results reported using scalp EEG based biometric systems [10], [11], never explored with this spatial resolution, further suggesting that these systems, other than show a very high uniqueness, may provide very good permanence properties. On the other hand, these results also confirm what is generally observable by designing a brain computer interface system. In fact, even though it is still remarkable a strong effect of task-switching, it is still evident that the individual traits may strongly hinder the generalization of the approach (failing to keep a good performance across different subjects).

## V. CONCLUSION

In conclusion, we have shown that source-level EEG analysis confirms that PSD, PLV and PLV derived functional brain network, as measured by nodal centrality (namely, eigenvector centrality), are stable over-time, dominated by individual properties but largely dependent from the specific task. These findings may have important implications for both clinical (e.g., biomarkers) and bio-engineering applications (e.g., biometric systems and brain computer interfaces).

## REFERENCES

[1] O. Miranda-Dominguez et al., “Connectotyping: Model Based Fingerprinting of the Functional Connectome,” PLOS ONE, vol. 9, no. 11, p. e111048, Nov. 2014.

[2] E. S. Finn et al., “Functional connectome fingerprinting: identifying individuals using patterns of brain connectivity,” Nature Neuroscience, vol. 18, no. 11, pp. 1664–1671, Nov. 2015.

[3] E. S. Finn and R. Todd Constable, “Individual variation in functional brain connectivity: implications for personalized approaches to psychiatric disease,” Dialogues Clin Neurosci, vol. 18, no. 3, pp. 277–287, Sep. 2016.

[4] M. Arns, “EEG-Based Personalized Medicine in ADHD: Individual Alpha Peak Frequency as an Endophenotype Associated with Nonresponse,” Journal of Neurotherapy, vol. 16, no. 2, pp. 123–141, Apr. 2012.

[5] C. Horien, X. Shen, D. Scheinost, and R. T. Constable, “The individual functional connectome is unique and stable over months to years,” NeuroImage, vol. 189, pp. 676–687, Apr. 2019.

[6] O. Miranda-Dominguez, E. Feczko, D. S. Grayson, H. Walum, J. T. Nigg, and D. A. Fair, “Heritability of the human connectome: A connectotyping study,” Network Neuroscience, vol. 2, no. 2, pp. 175–199, Nov. 2017.

[7] M. Demuru et al., “Functional and effective whole brain connectivity using magnetoencephalography to identify monozygotic twin pairs,” Scientific Reports, vol. 7, no. 1, p. 9685, Aug. 2017.

[8] M. DelPozo-Banos, C. M. Travieso, C. T. Weidemann, and J. B. Alonso, “EEG biometric identification: a thorough exploration of the time-frequency domain,” J Neural Eng, vol. 12, no. 5, p. 056019, Oct. 2015.

[9] M. Fraschini, A. Hillebrand, M. Demuru, L. Didaci, and G. L. Marcialis, “An EEG-Based Biometric System Using Eigenvector Centrality in Resting State Brain Networks,” IEEE Signal Processing Letters, vol. 22, no. 6, pp. 666–670, Jun. 2015.

[10] D. L. Rocca et al., “Human Brain Distinctiveness Based on EEG Spectral Coherence Connectivity,” IEEE Transactions on Biomedical Engineering, vol. 61, no. 9, pp. 2406–2412, Sep. 2014.

[11] M. Fraschini, S. M. Pani, L. Didaci, and G. L. Marcialis, “Robustness of functional connectivity metrics for EEG-based personal identification over task-induced intra-class and inter-class variations,” Pattern Recognition Letters, vol. 125, pp. 49–54, Jul. 2019.

[12] C. Gratton et al., “Functional Brain Networks Are Dominated by Stable Group and Individual Factors, Not Cognitive or Daily Variation,” Neuron, vol. 98, no. 2, pp. 439–452.e5, Apr. 2018.

[13] R. Cox, A. C. Schapiro, and R. Stickgold, “Variability and stability of large-scale cortical oscillation patterns,” Network Neuroscience, vol. 2, no. 4, pp. 481–512, Feb. 2018.

[14] M. Lai, M. Demuru, A. Hillebrand, and M. Fraschini, “A comparison between scalp- and source-reconstructed EEG networks,” Scientific Reports, vol. 8, no. 1, p. 12269, Aug. 2018.

[15] J.-P. Lachaux, E. Rodriguez, J. Martinerie, and F. J. Varela, “Measuring phase synchrony in brain signals,” Human Brain Mapping, vol. 8, no. 4, pp. 194–208, Jan. 1999.

[16] M. S. Hämäläinen and R. J. Ilmoniemi, “Interpreting magnetic fields of the brain: minimum norm estimates,” Med. Biol. Eng. Comput., vol. 32, no. 1, pp. 35–42, Jan. 1994.

[17] M. Hassan, O. Dufor, I. Merlet, C. Berrou, and F. Wendling, “EEG Source Connectivity Analysis: From Dense Array Recordings to Brain Networks,” PLoS One, vol. 9, no. 8, Aug. 2014.

[18] M. Demuru, S. M. L. Cava, S. M. Pani, and M. Fraschini, “A comparison between power spectral density and network metrics: an EEG study,” Apr. 2019.

[19] A. Delorme and S. Makeig, “EEGLAB: an open source toolbox for analysis of single-trial EEG dynamics including independent component analysis,” J. Neurosci. Methods, vol. 134, no. 1, pp. 9–21, Mar. 2004.

[20] F. Tadel, S. Baillet, J. C. Mosher, D. Pantazis, and R. M. Leahy, “Brainstorm: a user-friendly application for MEG/EEG analysis,” Comput Intell Neurosci, vol. 2011, p. 879716, 2011.

[21] A. Gramfort, T. Papadopoulo, E. Olivi, and M. Clerc, “OpenMEEG: opensource software for quasistatic bioelectromagnetics,” Biomed Eng Online, vol. 9, p. 45, Sep. 2010.

[22] F.-H. Lin, T. Witzel, S. P. Ahlfors, S. M. Stufflebeam, J. W. Belliveau, and M. S. Hämäläinen, “Assessing and improving the spatial accuracy in MEG source localization by depth-weighted minimum-norm estimates,” NeuroImage, vol. 31, no. 1, pp. 160–171, May 2006.

[23] R. S. Desikan et al., “An automated labeling system for subdividing the human cerebral cortex on MRI scans into gyral based regions of interest,” NeuroImage, vol. 31, no. 3, pp. 968–980, Jul. 2006.

[24] M. Fraschini, M. Demuru, A. Crobe, F. Marrosu, C. J. Stam, and A. Hillebrand, “The effect of epoch length on estimated EEG functional connectivity and brain network organisation,” J Neural Eng, vol. 13, no. 3, p. 036015, 2016.

[25] B. Ruhnau, “Eigenvector-centrality — a node-centrality?,” Social Networks, vol. 22, no. 4, pp. 357–365, Oct. 2000.

[26] Y. Benjamini, A. M. Krieger, and D. Yekutieli, “Adaptive linear step-up procedures that control the false discovery rate,” Biometrika, vol. 93, no. 3, pp. 491–507, Sep. 2006.

